# Collective unstructured interactions drive chromatin binding of transcription factors

**DOI:** 10.1101/2025.05.16.654615

**Authors:** Abrar A Abidi, Gina M Dailey, Robert Tjian, Thomas G W Graham

## Abstract

Eukaryotic transcription factors (TFs) contain both structured DNA-binding domains (DBDs) and intrinsically disordered regions (IDRs). While the structures and sequence preferences of DBDs have been extensively characterized, the role of IDR-mediated interactions in chromatin binding and nuclear organization remains poorly understood, in part because these interactions have been difficult to measure in living cells. Here, we use a recently developed single-molecule technique, proximity-assisted photoactivation (PAPA), to investigate how IDRs influence TF associations with each other and with chromatin, focusing on the factors Sp1 and Klf1. We find that the number and patterning of aromatic and basic residues within IDRs govern both TF self-association and chromatin binding. Unexpectedly, the isolated DBD of Sp1 binds chromatin very weakly and non-specifically. The isolated IDR, by contrast, interacts poorly with chromatin-bound wild-type Sp1, yet this interaction is enhanced when even minimal DNA-binding capacity is restored. Strikingly, replacing Sp1’s native DBD with those of heterologous TFs recovers both IDR-mediated interactions and chromatin association, despite divergent sequence preferences. PAPA measurements also reveal extensive heterotypic interactions between wild-type Sp1 and other TFs. Together, these results establish PAPA as a powerful method for studying unstructured interactions in their native context and suggest that IDRs participate in widespread cooperative associations scaffolded by transient DBD-DNA contacts, which concentrate disordered regions along chromatin. In contrast to classical models, we propose that TF specificity *in vivo* emerges not solely from DBD sequence preferences, but from a constellation of weak, dynamic, and diverse interactions mediated by IDRs.

## Introduction

For decades, the prevailing model for transcription factor (TF) specificity has maintained that structured DNA-binding domains (DBDs) locate their targets by recognizing well-defined sequence motifs along the genome (Brent & Ptashne, 1985; Stormo, 2013; Lambert et al., 2018). This framework—rooted in structural biology, *in vitro* biochemistry, and genomics—has grown increasingly misaligned with experimental evidence (Biggin, 2011; Mahendrawada et al., 2025). Although eukaryotic genomes are replete with potential binding sites for any given TF, only a small fraction are occupied *in vivo* (Tanay, 2006; Lambert et al., 2018; Horton et al., 2023). Moreover, different TFs that recognize the same motif often occupy distinct genomic loci within the same cell type (Hollenhorst et al., 2009; Brodsky et al., 2020; Gera et al., 2022), and many occupied sites lack recognizable motifs altogether (Slattery et al., 2014). These contradictions call into question whether DBD sequence preferences are sufficient—or even primary—determinants of genomic localization. In this study, we confront the paradox of how TFs reliably achieve specificity in a nuclear environment where potential binding sites are both overabundant and underspecified.

Unlike their bacterial counterparts, eukaryotic TFs contain extensive intrinsically disordered regions (IDRs)—polypeptide segments that lack stable three-dimensional structure (Sigler, 1988). Recent studies suggest that IDRs influence the genomic localization of TFs (Brodsky et al., 2020; Chen et al., 2022), yet the molecular basis of this contribution remains poorly understood. The distinctive properties of IDRs may be central to resolving this question, yet their behavior in living cells has remained largely inaccessible. A major obstacle has been methodological, with no reliable tools available to measure the weak, transient interactions characteristic of unstructured proteins in their true cellular context.

Prior studies have relied heavily on ectopic overexpression, artificial multimerization, or recruitment to synthetic gene arrays to visualize co-localization at bulk resolution (Boija et al., 2018; Wei et al., 2020; Chong et al., 2018). By inducing the formation of large, synthetic nuclear foci rather than resolving individual molecular interactions, these approaches often distort the very behaviors they seek to capture (McSwiggen et al., 2019). This has left most models of nuclear IDR function grounded in observations from highly engineered or non-physiological systems, rather than direct measurements of endogenous TFs under native conditions.

We overcame this technical barrier with proximity-assisted photoactivation (PAPA), a fluorescence imaging strategy recently developed to detect protein interactions *in vivo* at single-molecule resolution. By measuring spatial proximity, PAPA is uniquely suited to capture low-affinity, transient interactions that have evaded conventional methods (Graham et al., 2022; Graham et al., 2024; Dahal et al., 2025). To extend these measurements across a broad range of interactions, we built a custom oblique line-scan (OLS) microscope, which synchronizes a swept light sheet with the high-speed readout of an sCMOS camera (Driouchi, 2025).

We used PAPA to investigate the unstructured interactions of TFs, focusing on the human TFs Sp1 and Klf1 as case studies. We found that IDRs drive TF co-localization on chromatin through multivalent self-association. These interactions depend on both the number and patterning of aromatic residues within IDRs, and are further stabilized by basic residues in unstructured regions flanking the DBD, likely through electrostatic contacts with nucleic acids. Strikingly, the isolated Sp1 DBD bound chromatin poorly in cells and did not co-localize with the full-length protein.

In contrast, the isolated Sp1 IDR, lacking a DBD, still exhibited attenuated interactions with wild-type Sp1. Notably, fusing the IDR to heterologous DBDs—regardless of their sequence preferences—significantly enhanced both chromatin binding and interaction with the full-length protein. Even fusing Sp1’s DBD to IDRs from functionally unrelated RNA-binding proteins increased chromatin association and restored co-localization with wild-type Sp1, indicating that heterotypic interactions can also contribute to genomic localization. Finally, we observed widespread heterotypic interactions between Sp1 and other TFs, consistent with a model in which a broad repertoire of unstructured interactions collectively shapes TF occupancy across the genome.

Taken together, these findings suggest that individual DBDs are not the sole—or even primary— determinants of TF specificity, but rather one component of a higher-order regulatory architecture in which IDRs play a central role. The insufficiency and relative nonspecificity of eukaryotic DBDs may reflect a deeper evolutionary shift: as regulatory complexity increased, rigid, deterministic sequence recognition gave way to a more flexible and integrative mode of gene regulation. Synthesizing our observations, we propose a model of *emergent specificity*, in which genomic localization arises from the combined contributions of DBDs and IDRs, acting through ensembles of weak, preferential interactions concentrated along chromatin.

## Results

### PAPA Detects Weak Unstructured Interactions in Living Cells

Proximity-assisted photoactivation (PAPA) is an optical technique we use to detect protein-protein interactions at single-molecule resolution in live cells. It relies on a photoswitching behavior observed with Janelia Fluor dyes: two proteins of interest are fused to self-labeling tags, Halo and SNAPf, which are labeled with “sender” (JF549) and “receiver” (JFX650) fluorophores, respectively (Fig. 1A). The receiver fluorophore, JFX650, is first converted to a nonfluorescent dark state by red (642 nm) light. A brief pulse of green (560 nm) light then excites the JF549 sender, selectively reactivating receiver-labeled proteins that are in close proximity to the sender. These reactivated molecules can then be tracked through single-molecule imaging. As an internal control, violet (405 nm) light triggers proximity-independent direct reactivation (DR), providing a sample of the total population of receiver-labeled molecules.

**Figure 1.**
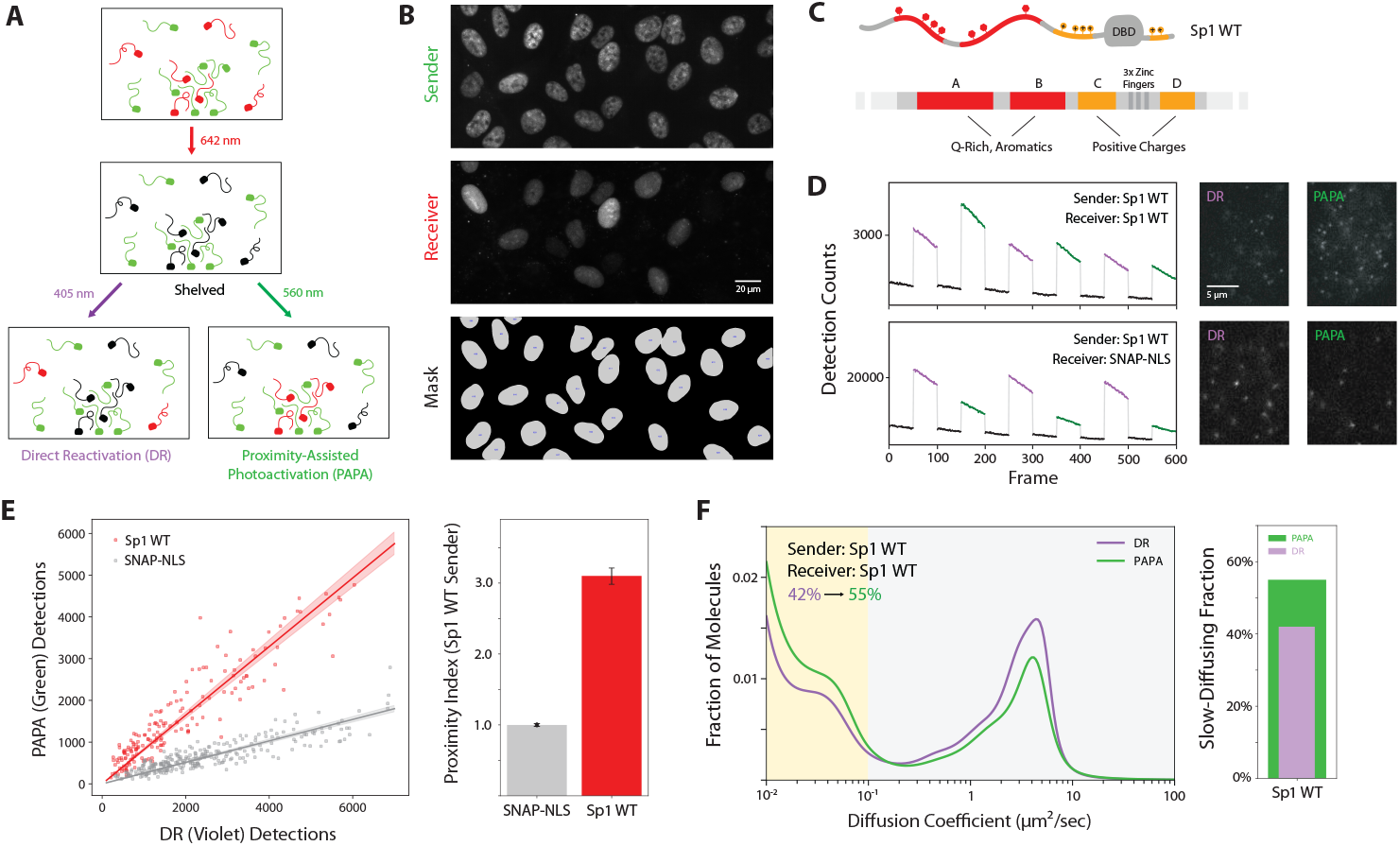
PAPA detects Sp1 self-association in living cells. **(A)** Schematic of proximity-assisted photoactivation (PAPA). Receiver-labeled proteins (red) are shelved into a nonfluorescent dark state using 642 nm light. Subsequent excitation of nearby sender-labeled proteins (green) with 560 nm light selectively reactivates nearby receivers. As a control, 405 nm illumination induces direct reactivation (DR) of receivers independent of their proximity to the sender. **(B)** High-throughput single-molecule imaging using an OLS microscope. Representative widefield images show sender (JF549-Halo-Sp1), receiver (JFX650-SNAPf-Sp1), and nuclear segmentation masks used for quantification. **(C)** Domain diagram of Sp1. The N-terminal A and B regions are enriched in glutamine and aromatic residues, while the C and D regions flanking the DBD are enriched in basic residues. **(D)** Detection counts over time under alternating violet (DR) and green (PAPA) illumination for cells co-expressing Sp1 as both sender and receiver (top) or Sp1 with a non-interacting nuclear SNAPf control (bottom). Insets show representative single-molecule frames for each condition and illumination mode. **(E)** Left: Scatterplot of DR versus PAPA detection counts per nucleus. Right: Bar plot showing proximity index (PAPA/DR), normalized to non-interacting SNAPf-NLS control. **(F)** Left: Diffusion coefficient distributions of PAPA- and DR-reactivated molecules. Right: Slow-moving (< 0.1 µm^2^/s) fraction of molecules reactivated by PAPA (green) and DR (violet).

To increase the throughput of these measurements, we built an OLS microscope, which sweeps a light sheet over the specimen in synchrony with the readout of an sCMOS camera (Driouchi, 2025). This provides a much wider field of view than standard HILO microscopy (Fig. 1B) while maintaining a fast frame rate and high signal-to-noise. Because our previous applications of PAPA involved more stable structured interactions, it was unknown whether PAPA could resolve weaker, transient interactions such as those mediated by IDRs. To test this, we applied PAPA to the TF Sp1 (Fig. 1C).

We used Cas9 genome editing to insert HaloTag at the endogenous *Sp1* locus in U2OS cells and simultaneously expressed a SNAPf-Sp1 fusion protein from a doxycycline-inducible transgene integrated at the *AAVS1* “safe harbor” locus (Fig. S1A-B). This strategy allowed precise control over copy number and expression level of Sp1 variants. Flow cytometry confirmed that transgene expression remained similar to or below endogenous levels, minimizing artifacts associated with overexpression (Fig. S1C).

For each condition, we alternated imaging cycles with green (PAPA) and violet (DR) excitation and counted the number of reactivated receiver molecules (Fig. 1D, E). These counts were compared to cells expressing nuclear SNAPf alone as a non-interacting control. We define a “proximity index” as the ratio of PAPA to DR reactivation events, normalized to this control (Fig. 1E). A proximity index greater than one indicates that the proteins are in close proximity (within ∼20 nm) more often than expected by chance.

PAPA revealed robust self-association between Sp1 molecules in live cells (Fig. 1E). State array analysis of PAPA detections also showed an enrichment of molecules with the slow mobility characteristic of chromatin-bound proteins (Fig. 1F), suggesting that Sp1 self-association correlates with chromatin localization. We refer to this population as “chromatin-bound” for brevity, although we discuss potential caveats to this interpretation in the Discussion.

We applied the same approach to Klf1, a related zinc-finger TF that is not natively expressed in U2OS cells. Using a bicistronic transgene inserted at the *AAVS1* locus, we co-expressed combinations of either Halo-Klf1 and SNAPf-Klf1 or Halo-Klf1 and a nuclear SNAPf control. As with Sp1, PAPA detected robust self-association of Klf1 relative to the non-interacting control (Fig. S2A).

### Aromatic Residues Promote Self-Association on Chromatin

We next investigated the role of IDRs in Sp1 self-interaction. Sp1’s N-terminal IDR contains two glutamine- and aromatic-rich segments, labeled A and B (Fig. 1C). Using the inducible transgene system described above, we assayed interaction of SNAPf-tagged deletion mutants with endogenous Halo-tagged Sp1 (Fig. 2A). Both A and B deletion mutants showed less PAPA signal with full-length Sp1 and reduced chromatin interaction (Fig. 2B-C). Nonetheless, PAPA detected enriched chromatin-bound molecules, indicating that association of the mutants with full-length Sp1 correlated with chromatin binding (Fig. 2C). As expected, the ΔA+B double mutant was more severely impaired in full-length Sp1 interaction and chromatin association, suggesting that the A and B regions contribute additively to self-association (Fig. 2B-C).

**Figure 2.**
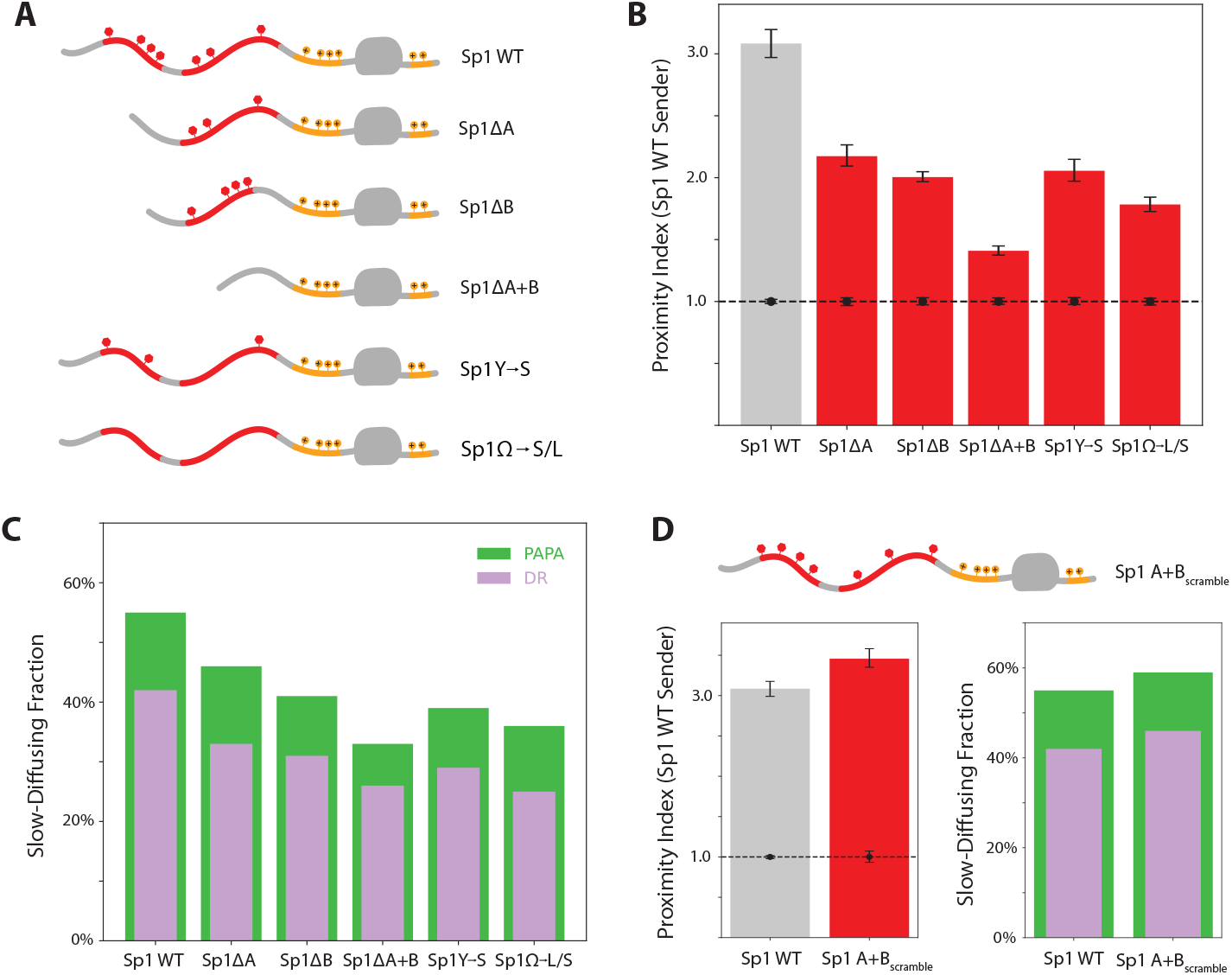
Aromatic residues in Sp1 IDRs drive self-association and chromatin binding. **(A)** Diagrams of aromatic-enriched region mutants. **(B)** PAPA signal between full-length Sp1 and each variant in (A). A dashed line at proximity index 1 represents the non-interacting control imaged for that session, with its associated error bars. **(C)** Slow-diffusing (< 0.1 µm^2^/s) fraction of molecules reactivated by PAPA (green) and DR (violet). **(D)** Left: Effect of the A+B_scramble_ variant on PAPA signal with full-length Sp1. Right: Corresponding slow-diffusing fractions.

To assess the role of aromatic residues and rule out indirect effects from changes in protein length, we generated variants in which either all tyrosine (Sp1Y→S) or all aromatic residues (Sp1Ω→S/L) outside the DBD were selectively mutated to non-aromatic residues (Fig. 2A). Both variants showed substantially reduced association with full-length Sp1 and with chromatin (Fig. 2B-C). The no-tyrosine mutant produced a milder phenotype than the no-aromatic mutant, implying that self-association scales with total aromatic content.

To test whether the amino acid sequence, not just the total aromatic content, influenced self-association, we scrambled the amino acid sequences of segments A and B while preserving their overall composition (Fig. 2D). This redistribution dispersed the aromatic residues, reducing their local clustering—an alteration that, based on prior simulations and *in vitro* studies, was expected to weaken self-association (Martin et al., 2020). Surprisingly, the scrambled variant exhibited a higher proximity index than the wild-type protein and showed a slightly increased fraction of chromatin-associated molecules (Fig. 2D). These results indicate that the Sp1 IDR not only tolerates sequence reordering, but can exhibit stronger self-association as a result.

The effect of residue arrangement on Sp1’s self-association prompted us to ask whether naturally occurring sequence variation might have similar effects. While Sp1 is highly conserved across mammals—showing 96% sequence identity between human and mouse—Klf1 exhibits greater divergence, with a more modest 71% identity, largely due to differences in the N-terminal disordered region. Although the total number of aromatic residues is preserved, their identities and spacing differ between species. Strikingly, mouse Klf1 exhibited a slightly higher proximity index with human Klf1 than human Klf1 with itself, suggesting that evolutionary shifts in residue order have influenced the interactions of the Klf1 IDR (Fig. S2B).

Taken together, these results indicate that Sp1 and Klf1 self-interactions in live cells are strongly dependent on their IDRs and influenced by the number and positioning of aromatic residues.

### Basic Residues Flanking the DBD Stabilize Chromatin Binding

Sp1 contains two positively charged unstructured regions flanking its DBD: the N-terminal C region and the C-terminal D region (Fig. 1C). Such basic stretches are common among eukaryotic TFs (Oksuz et al., 2023), yet their role in chromatin engagement remains poorly understood. To investigate how these basic regions might influence Sp1 interactions, we generated single and double deletion mutants lacking the C and D regions (Fig. 3A). Both Sp1ΔC and Sp1ΔD exhibited reduced interaction with endogenous Sp1, and the double mutant Sp1ΔC+D showed a further decrease (Fig. 3B). These interaction defects were milder than those observed for deletions of the aromatic-rich A and B segments (Fig. 2B). In contrast, chromatin association was more strongly impaired in the C and D deletion variants than in the A and B deletions (Fig. 2C & 3C). These findings suggest that the basic residues flanking the DBD may stabilize chromatin binding—likely through electrostatic interactions with negatively charged nucleic acids. Destabilizing chromatin association, in turn, appears to weaken Sp1 self-association, underscoring that these two types of interaction are likely interdependent.

**Figure 3.**
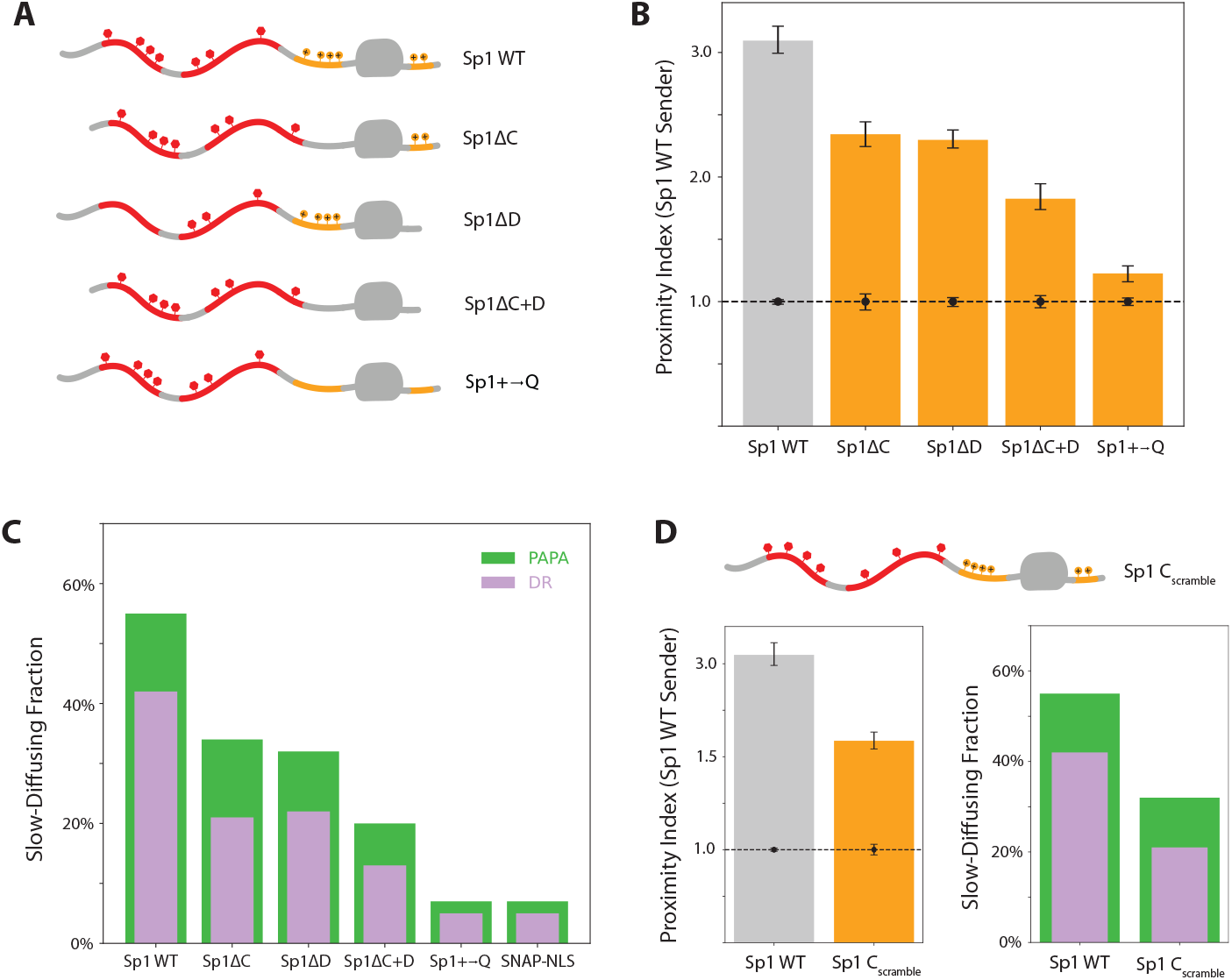
Basic residues flanking the Sp1 DBD stabilize self-association and are necessary for chromatin binding. **(A)** Diagrams of basic-enriched region variants. **(B)** PAPA signal between full-length Sp1 and each variant in (A). **(C)** Slow-diffusing (< 0.1 µm^2^/s) fraction of molecules reactivated by PAPA (green) and DR (violet). **(D)** Left: Effect of the C_scramble_ variant on PAPA with full-length Sp1. Right: Corresponding slow-diffusing fractions.

To rule out confounding effects from changes in protein length and to neutralize positive charge beyond the C and D regions, we generated a variant in which all lysine, arginine, and histidine residues outside the DBD were mutated to glutamine (Sp1+→Q; Fig. 3A). Strikingly, this variant exhibited minimal interaction with wild-type Sp1 and showed almost no chromatin association (Fig. 3B-C).

We initially reasoned that the precise positioning of basic residues might not be critical, given that flexible, unstructured regions can adopt a wide range of conformations. However, scrambling the C region, which increased the average distance of basic residues from the DBD, reduced both self-association and chromatin binding (Fig. 3D). These results suggest that the stabilizing effect of basic residues may depend not only on their presence within the IDR but also on their proximity to the DBD.

### The Core Sp1 DBD is Insufficient for Chromatin Binding

The basic-rich regions flanking Sp1’s DBD lack the structural rigidity and stereospecificity typically associated with sequence-specific nucleic acid binding domains. Yet in their absence, Sp1 loses its ability to stably bind chromatin, despite retaining an intact DBD and the full complement of aromatic-rich disordered regions. To test whether the loss of chromatin binding resulted from an indirect disruption of DBD function caused by truncating the basic-rich regions, we examined the isolated DBD.

Previous DNase footprinting experiments showed that the isolated DBD of Sp1 readily binds GC box sequences *in vitro* (Kadonaga et al., 1988). Clustered GC box motifs are abundant throughout the human genome, particularly within CpG island promoters and other GC-rich regulatory regions, where they often occur in dense arrays upstream of transcription start sites (Gardiner-Garden and Frommer, 1987; Yang et al., 2007). Given this fact, one might expect that the isolated DBD would show co-localization with wild-type Sp1 molecules bound to nearby recognition sequences.

Unexpectedly, we found that the isolated DBD of Sp1 exhibited very low chromatin association (Fig. 4A) and negligible PAPA signal with endogenous Sp1 (Fig. 4B, S3). Similar proximity indices were observed between endogenous Sp1 and histone H2B, a core constituent of chromatin that is not known to interact directly with Sp1 (Fig. 4B).

**Figure 4.**
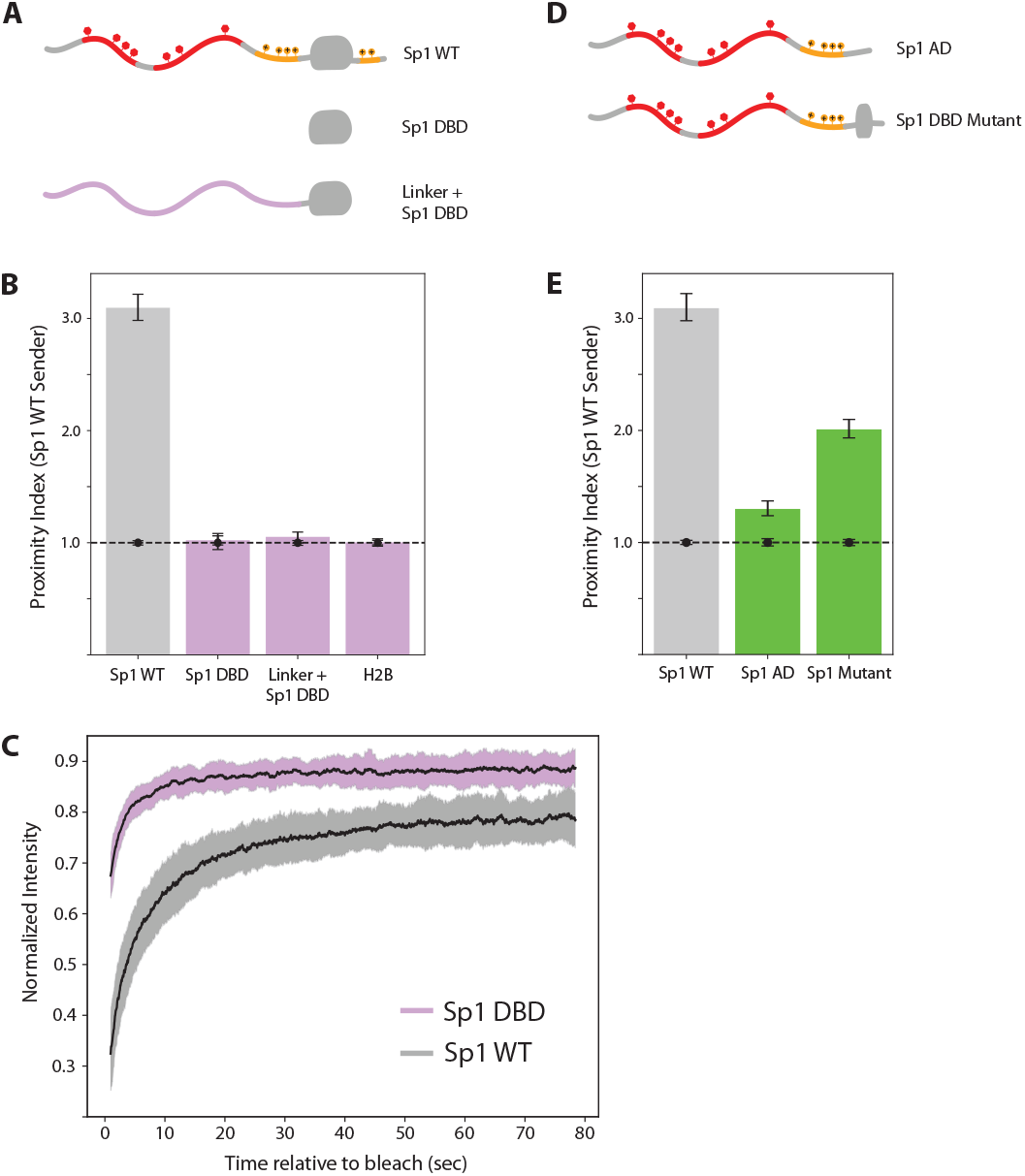
Figure 4. Live imaging of the isolated Sp1 DBD and IDR. **(A)** Diagrams of IDR deletion and linker replacement mutants. **(B)** PAPA signal between full-length Sp1 and each variant in (A), with histone H2B-SNAPf as an additional non-interacting negative control. **(C)** Fluorescence recovery after photobleaching (FRAP) curves comparing full-length Sp1 and the isolated DBD. **(D)** Diagrams of the isolated Nterminal IDR (Sp1 AD) and single-zinc finger mutant (Sp1 DBD Mutant). **(E)** PAPA signal between full-length Sp1 and the variants in (D).

To rule out the possibility that the failure to detect interactions stemmed from steric hindrance by the SNAPf tag or from changes in inter-dye distance affecting PAPA efficiency, we replaced Sp1’s N-terminal IDR with a synthetic flexible linker of equivalent length. This variant also failed to interact with wild-type Sp1 and showed negligible chromatin association, confirming that the DBD alone is insufficient for self-interaction or stable chromatin binding, irrespective of the fluorophore’s distance from the DBD (Fig. 4B, S3). Supporting this conclusion, FRAP analysis revealed a markedly shallower bleach depth and faster fluorescence recovery for the isolated DBD compared to full-length Sp1 (Fig. 4C).

These results show, contrary to expectations rooted in *in vitro* DNA binding studies, that the structured DBD of Sp1 binds poorly to chromatin in cells and fails to co-localize with full-length Sp1 in the absence of its IDRs.

### Weak DNA Binding Concentrates IDRs Along Chromatin

Given that the N-terminal IDR of Sp1 is necessary for chromatin binding and interaction with full-length Sp1 in cells, we next asked if it is sufficient. An N-terminal IDR fragment showed a weak though measurable PAPA signal with full-length Sp1 (Fig. 4E, middle bar). A similarly weak but measurable interaction was likewise detected between full-length Klf1 and Klf1’s IDR (Fig. S2C).

We next examined a disease-associated Sp1 mutant containing a heterozygous frameshift that preserves the N-terminal IDR but retains only a single zinc finger in the DBD (Tummala et al., 2020). DNA binding by this truncated DBD, if any, is predicted to be weak and nonspecific, and we thus expected that its behavior would resemble that of the IDR alone. Remarkably, this Zn-finger mutant showed significantly stronger interaction with wild-type Sp1 than the isolated IDR, suggesting that even weak, nonspecific DNA binding can stabilize IDR-mediated interactions between TFs.

To test the generality of this phenomenon, we examined the E325K mutation in Klf1, which causes a dominant form of congenital dyserythropoietic anemia type IV (Arnaud et al., 2010). This mutation substitutes a glutamate in the second zinc finger with a basic residue and is thought to confer altered DNA-binding specificity (Varricchio et al., 2019). Similar to the Sp1 mutant, the Klf1 mutant retained substantial interaction with wild-type Klf1 (Fig. S2C), exceeding that of the isolated N-terminal IDR—underscoring that altered DBD sequence preferences do not necessarily disrupt preexisting IDR–IDR associations.

### DBD Sequence Preferences are Not Essential for TF Co-localization

To further investigate how the sequence-specificity of DBDs might influence unstructured interactions, we replaced Sp1’s DBD with two structurally and functionally distinct alternatives: the DBD of human CTCF and the bacterial LacI protein (Fig. 5A). The former recognizes highly specific endogenous sequences, while the latter lacks specific binding sites in the human genome. Strikingly, both chimeras retained substantial interaction with wild-type Sp1, while neither DBD alone exhibited detectable interaction (Fig. 5B). These results reinforce the idea that DNA binding—whether sequence-specific or not—can stabilize IDR-IDR interactions.

**Figure 5.**
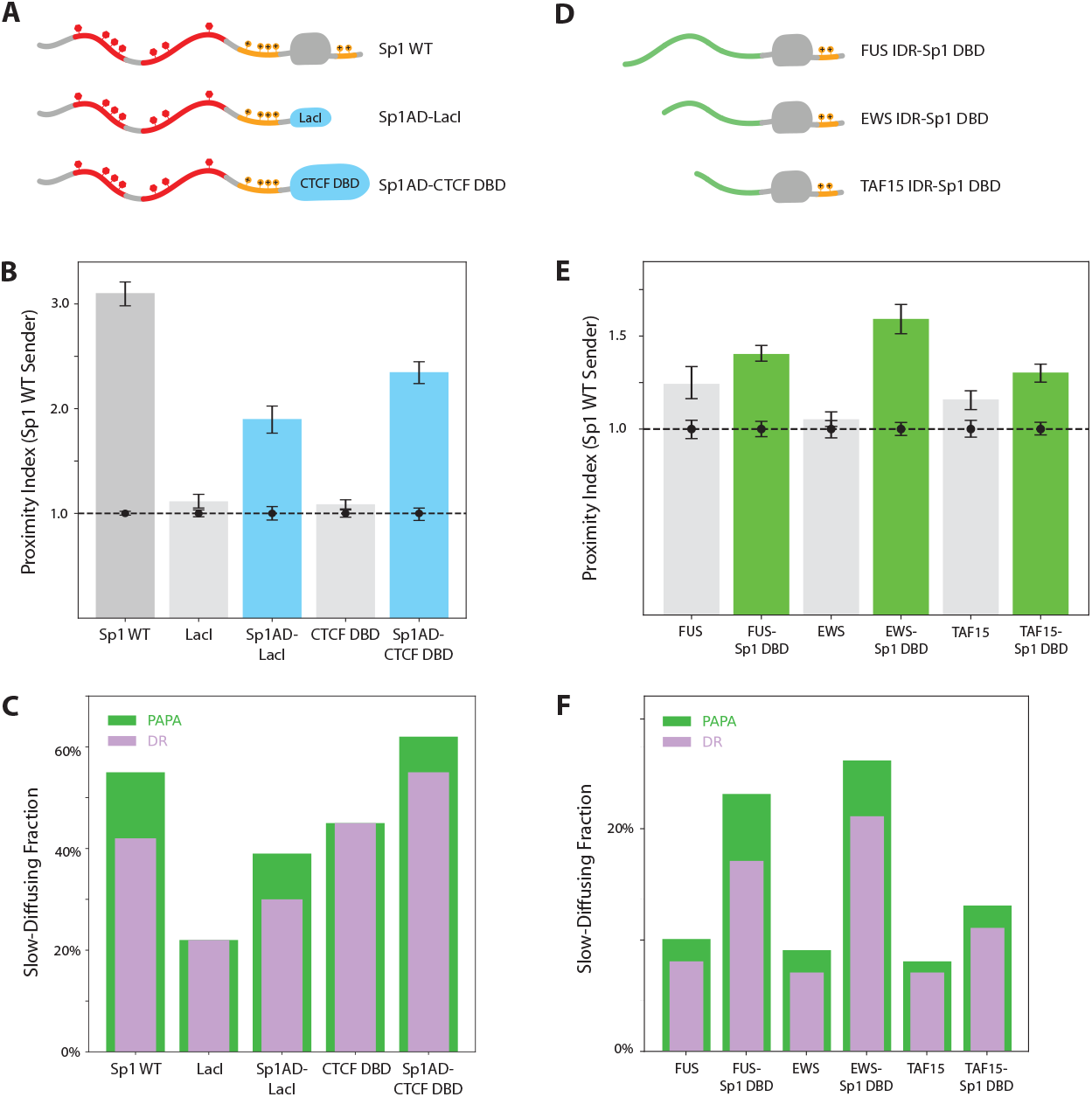
Sp1 variants with heterologous DBD or IDR swaps retain association with wild-type Sp1. (A) Diagrams of wild-type Sp1 and chimeric variants in which the Sp1 DBD is replaced with LacI or the CTCF DBD. PAPA signal between full-length Sp1 and each variant in (A). **(C)** Slow-moving (< 0.1 µm^2^/s) fraction of molecules reactivated by PAPA (green) and DR (violet) for each variant in (A). **(D)** Diagrams of constructs in which IDRs from FUS, EWS, or TAF15 replace the N-terminal IDR of Sp1. **(E)** PAPA signal between full-length Sp1 and each variant in (D). **(F)** Slow-diffusing fractions of PAPA/DR-reactivated molecules for each variant in (D).

### Chromatin Binding Facilitates Unstructured Interactions with Heterologous TFs

To assess the generality of IDR-mediated TF-TF interactions, we first asked whether IDRs from three different RNA-binding proteins could associate with the IDR of Sp1. A previous study using a synthetic gene array found no significant co-localization between Sp1’s IDR and those of EWS, FUS, or TAF15 (Chong et al., 2018). Revisiting these interactions using PAPA in live cells under physiological conditions, we similarly observed little to no detectable interactions between the freely diffusing IDRs and wild-type Sp1 on native chromatin (Fig. 5E).

However, when these same IDRs were fused to DBDs (Fig. 5D), they exhibited increased interactions with Sp1 (Fig. 5E) and a higher proportion of chromatin-bound, slow-diffusing molecules (Fig. 5F). These results suggest that chromatin binding can expose latent interaction potential between IDRs that would otherwise interact poorly.

To test whether such heterotypic interactions reflect a broader feature of TF behavior, we measured PAPA signals between endogenous Sp1 and six other structurally distinct TFs, each possessing substantial IDRs (Fig. 6A). We observed a range of interactions above background, with PAPA selectively enriching chromatin-bound molecules (Fig. 6B). These findings suggest that Sp1 forms weak, chromatin-localized interactions with a surprisingly broad array of heterologous TFs.

**Figure 6.**
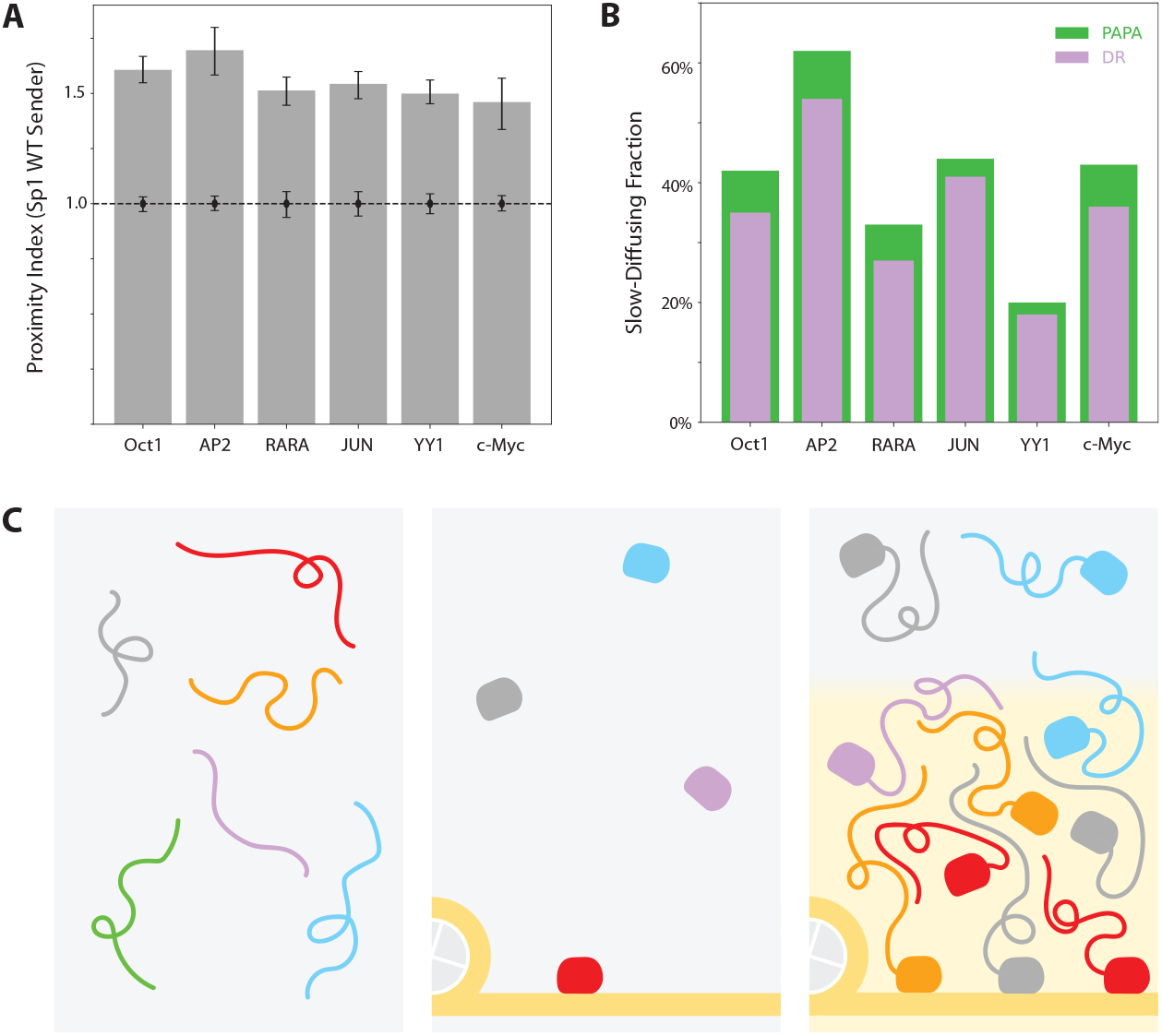
Sp1 shows extensive heterotypic interactions with other transcription factors. **(A)** PAPA signal between endogenous Sp1 and heterologous TFs. **(B)** Slow-diffusing (< 0.1 µm^2^/s) fraction of PAPA/ DR-reactivated molecules in (A). **(C)** Model of TF binding to chromatin. Left: Freely diffusing IDRs interact with low probability at endogenous concentrations. Middle: Isolated DNA-binding domains interact weakly and nonspecifically with chromatin. Right: IDR-mediated interactions between TFs, scaffolded by DBD-DNA interactions, promote assembly of TF ensembles.

## Discussion

The genomic localization of TFs is typically understood through a lock-and-key model, where structured DBDs recognize specific DNA sequence motifs positioned along the genome. This framework assumes a modular separation of function for TFs, with DBDs conferring DNA-binding specificity and IDRs driving transcriptional activation. Our findings challenge this view, with PAPA providing the key technical advance that allowed us to measure unstructured interactions between TFs in their native cellular context. Using a custom-built OLS microscope, we drastically increased the throughput of our PAPA measurements, allowing us to profile wild-type Sp1 interactions with an extensive panel of Sp1 variants in addition to other TFs.

We draw attention to three core observations from this study (Fig. 6C). First, DBDs alone showed very little chromatin binding and failed to co-localize with full-length TFs by PAPA. Second, TF IDRs showed measurable, but minimal, interaction with full-length TFs. Third—and most crucially—many different IDR-DBD fusions exhibited appreciable chromatin binding and interaction with full-length Sp1. This was true irrespective of DBD specificity and held, to a degree, even for IDRs from non-TFs.

Our findings reveal widespread cooperativity involving DBD-DNA and IDR-IDR interactions of TFs in live cells. Weak DBD–DNA contacts concentrate IDRs along chromatin, permitting interactions that otherwise occur infrequently in solution. Unstructured interactions with other TFs reciprocally stabilize even nonspecific DBD-DNA contacts.

Taken together, these observations suggest that the binding specificity of eukaryotic TFs is not an intrinsic property of individual DBDs, but a collective outcome of disordered interactions scaffolded by DBD-DNA contacts. Because this mode of regulation is irreducible to the intrinsic sequence preferences of individual domains, we refer to it as *emergent specificity*.

Enduring paradoxes—such as divergent binding profiles among TFs with near-identical DBDs, occupancy at sites lacking canonical motifs, and the pronounced influence of unstructured regions—may find a more coherent explanation under this framework. It may also help to explain why, across evolution, the sequence preferences of eukaryotic DBDs have become highly degenerate, while regions outside the DBD have expanded and acquired extensive disorder.

These collective interactions are qualitatively distinct from those characteristic of conventional models of TF cooperativity, which involve stereospecific interactions between folded domains positioned at fixed genomic sites. Historical examples like the interferon-β enhanceosome have strongly reinforced this view (Thanos & Maniatis, 1995; Panne et al., 2004). However, such models reflect the limitations of the tools that uncovered them—gel shifts, footprinting, and crystallography—which privilege structurally stable, high-affinity cooperative arrangements. In line with recent studies that have questioned the sufficiency of this framework (Vincent et al., 2016; Park et al., 2019; Kim et al., 2022; Mahendrawada et al., 2025), our live imaging results suggest a mode of cooperativity independent of either rigid DBD sequence preferences or structurally defined contacts between TFs.

Our findings may also have implications for how TF behavior is altered in disease. In the case of oncoproteins such as EWS-FLI1 (Delattre et al., 1994) and FUS-CHOP (Crozat et al., 1993), fusion of IDRs from the RNA-binding proteins EWS or FUS to structured DBDs is typically thought to preserve the DNA-binding preferences of the wild-type TF, driving pathological activation of its canonical genomic targets. Instead, our model suggests these oncoproteins may hijack the unstructured interactions of native TFs, altering their collective binding patterns. Many diseases are also linked to dominant heterozygous DBD mutations in TFs, including not only Sp1 and Klf1 but also PAX6, FOXP2, GATA3, and TCF4 (Azuma et al., 1998; Morison et al., 2023; Van Esch et al., 2000; Zweier et al., 2007). The disease-associated E325K mutation in Klf1, for example, is widely interpreted as redirecting genomic localization through altered motif specificity, consistent with a conventional view of TF binding (Ilsley, M.D. et al., 2019; Varricchio et al., 2019). However, we find that Sp1 and Klf1 DBD mutants retain co-localization with their wild-type counterparts, indicating that IDR-IDR interactions remain significant and may play a role in their pathology.

PAPA provides an unprecedented approach to measuring native IDR-mediated interactions in live cells, but it does not reveal the specific genomic loci where these interactions occur.

Moreover, single-molecule tracking cannot distinguish direct binding to DNA from indirect binding via intermediate molecules. Genomic-resolution methods such as ChIP and CUT&RUN could help fill these gaps, but their reliance on fixation and permeabilization may significantly alter the very unstructured interactions being investigated (Baranello, L. et al., 2016; Irgen-Gioro et al., 2022; Moroianu, J. & Blobel, G., 1995). A recent study linked increased chromatin-bound fractions of synthetic TFs to higher expression at a reporter locus (Fan et al., 2024), suggesting that chromatin association may correlate with transcriptional activity. Nevertheless, while our findings establish a strong link between IDR-IDR interactions and chromatin binding, further work is needed to determine how these interactions shape the transcriptional activity of endogenous genes in their native cellular context.

In short, our study reframes TF binding to chromatin as a property not of individual structured domains, but of dynamic, low-affinity interactions. These interactions, which we have visualized in live cells using PAPA, may represent a previously overlooked layer of transcriptional regulation with implications for evolution, development, and disease.

**Figure S1.**
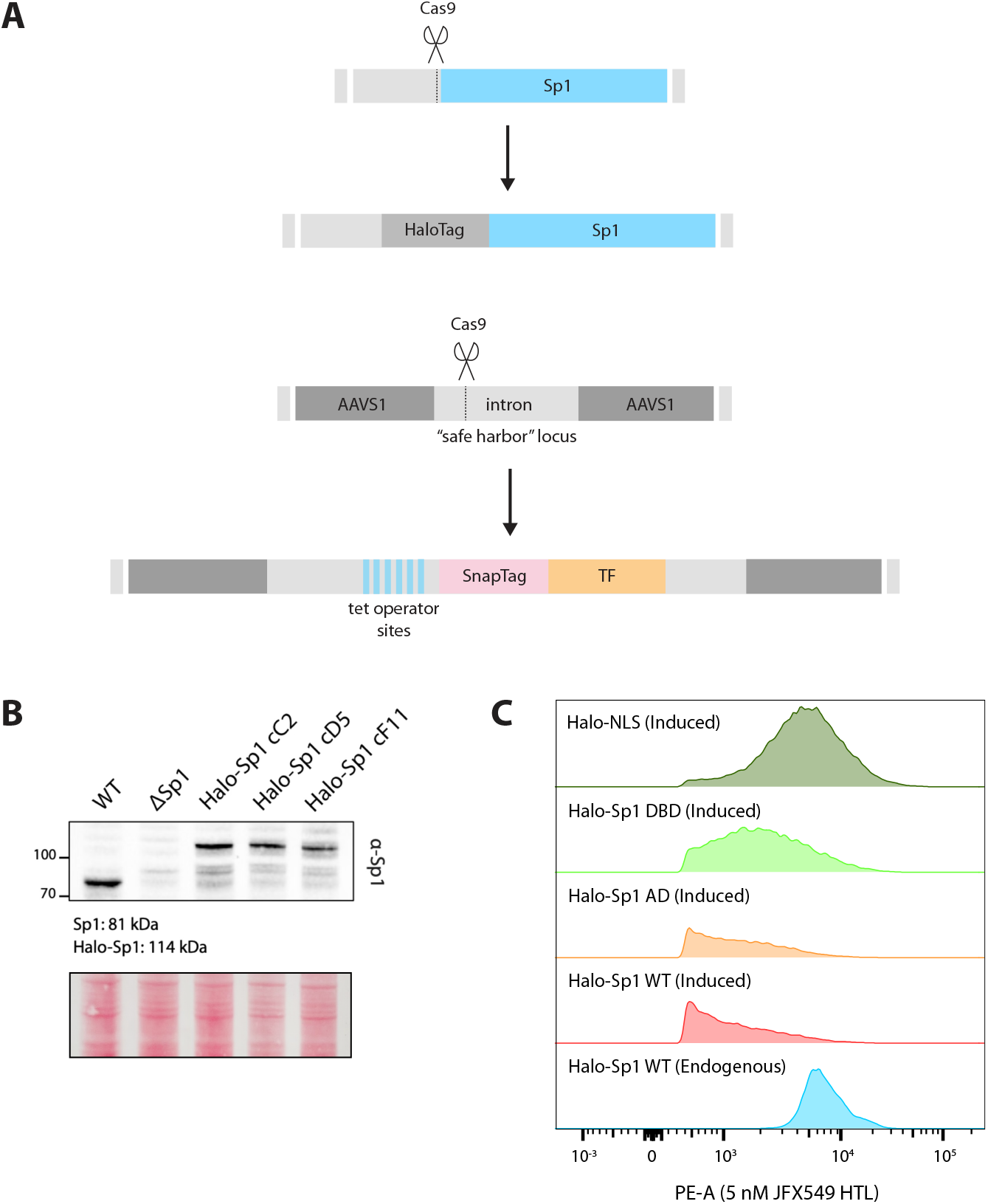
Engineering strategy for HaloTag and SNAPfTag fusions enables PAPA measurements at sub-physiological expression levels. **(A)** CRISPR-Cas9 editing was used to engineer a HaloTag fusion at the *Sp1* endogenous locus. SNAPfTag fusion proteins driven by an inducible promoter were integrated at the *AAVS1* locus. **(B)** Western blot shows correct band size for HaloTag knock-in as well as Sp1 knockout. **(C)** Flow cytometry quantifies fluorescence for inducible transgene expression and endogenous HaloTag-Sp1 fusion.

**Figure S2.**
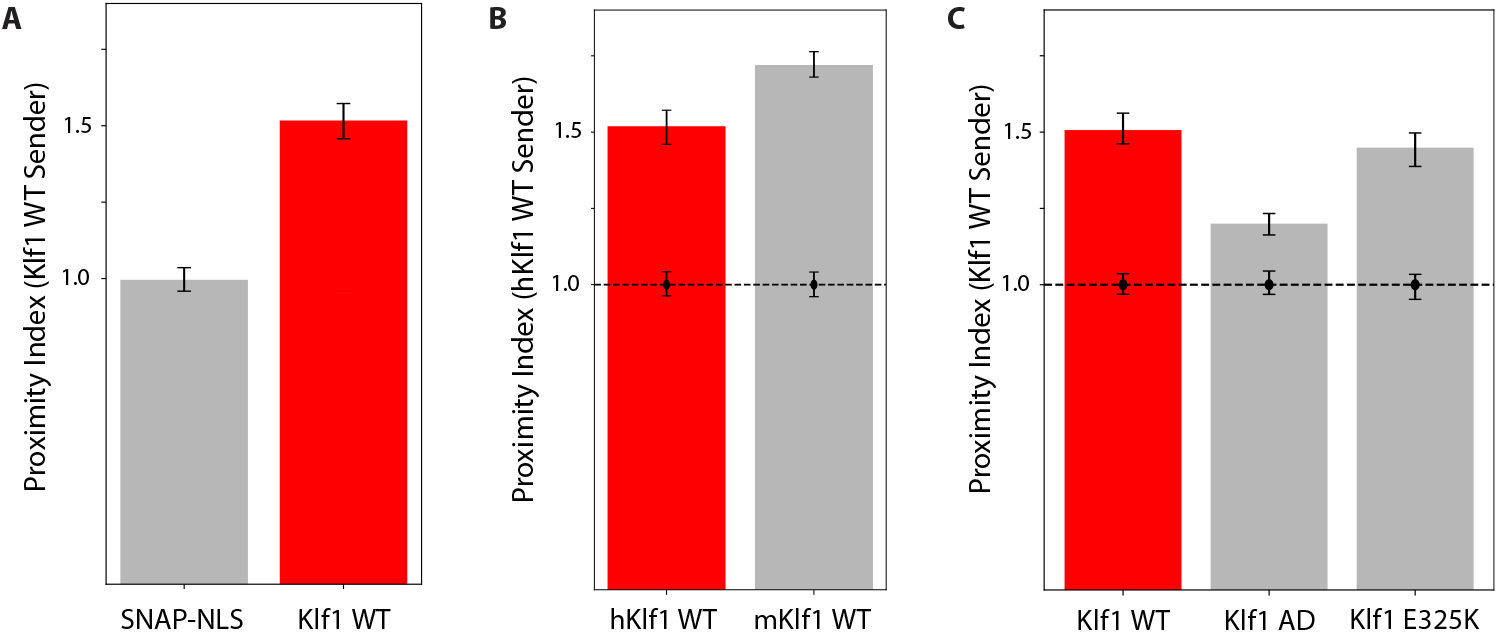
PAPA detects interactions between Klf1 and different Klf1 variants. **(A)** PAPA signal between sender-labeled and receiver-labeled wild-type Klf1. **(B)** PAPA signal between sender-labeled human Klf1 and receiver-labeled mouse Klf1. **(C)** PAPA signal between sender-labeled wild-type Klf1 and both receiver-labeled N-terminal Klf1 IDR (AD) and receiver-labeled E325K mutant Klf1.

**Figure S3.**
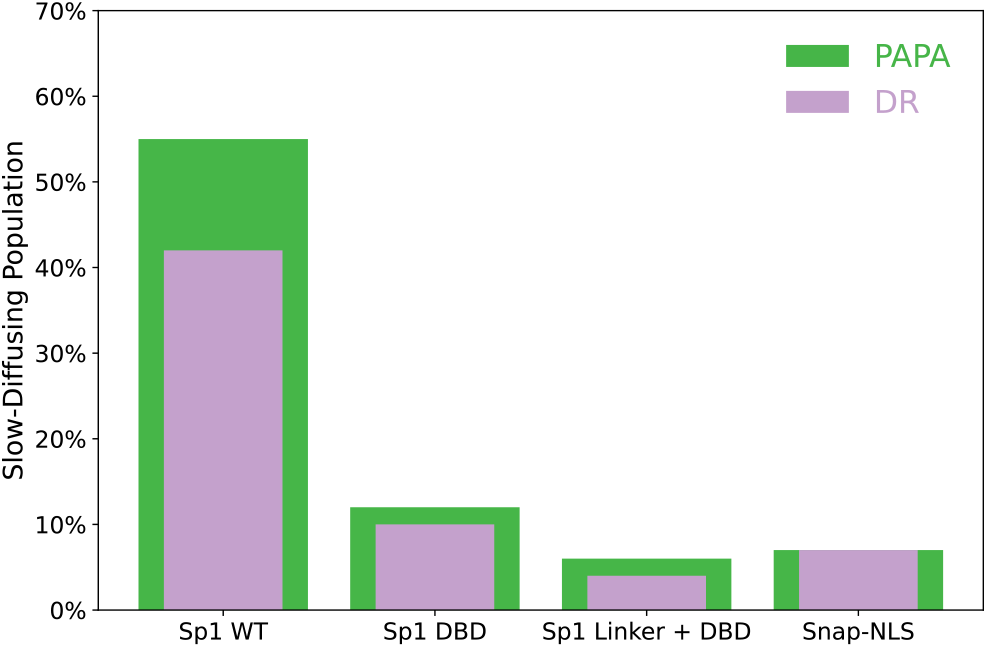
Isolated Sp1 DBD fails to appreciably bind chromatin.

## Methods

### Cell culture and cell line generation

U2OS cells were cultured at 37 °C with 5% CO_2_ in high-glucose Dulbecco’s Modified Eagle Medium (DMEM; Thermo Fisher #12800082) supplemented with 10% fetal bovine serum and penicillin/streptomycin. Cells were routinely tested for mycoplasma contamination by PCR.

To insert a HaloTag on endogenous *Sp1*, cells were co-transfected with a homology donor plasmid and a plasmid co-expressing Cas9 and an sgRNA. Cells were stained with a Janelia Fluor Halo ligand, and individual Halo-positive cells were flow-sorted into 96-well plates.

Homozygous knock-in clones were validated by PCR, western blot, and sequencing.

For *Sp1* knockout, cells were co-transfected with plasmids expressing Cas9 and sgRNAs targeting sequences flanking the *Sp1* coding region. Venus-positive cells (confirming construct delivery) were flow-sorted into 96-well plates, and deletion clones were validated by PCR and western blot.

Doxycycline-inducible transgenes were inserted into the *AAVS1* “safe harbor” locus using a homology-directed donor co-transfected with a plasmid expressing Cas9 and an sgRNA targeting *AAVS1*. The donor also carried a linked puromycin resistance cassette, allowing stable integrants to be selected in medium containing 1 µg/mL puromycin.

### Oblique line-scan microscopy

We constructed an oblique line-scan microscope on a Nikon Ti2 E inverted motorized microscope stand equipped with a SR Plan Apo IR 60X NA 1.27 water immersion objective lens. Laser lines (405 nm, 488 nm, 560 nm, and 642 nm) were combined using dichroic mirrors, directed through an acousto-optic tunable filter (AOTF), and passed through a single-mode optical fiber using two achromatic FiberPorts (ThorLabs PAF2-A7A). A light sheet was generated using a Powell lens (Laserline Optics, LOCP-8.9R10-1.2) and swept over the sample using a 1D galvo mirror (Thorlabs GVS201). A field-programmable gate array (FPGA) controlled by custom LabView software was used to modulate the AOTF and sweep the light sheet in synchrony with the exposure of a Hamamatsu Orca Fusion BT camera running in lightsheet mode. Samples were kept at 37 °C with 5% CO_2_ within an OKO Labs stage incubation chamber.

### PAPA-SMT

Cells were plated in 35-mm MatTek glass-bottom dishes in phenol-red free DMEM and induced for 36 hours with 1 µg/mL doxycycline while also staining with 50 nM of JF549 Halo ligand and 50 nM of JFX650 SNAP ligand. Prior to imaging, cells were washed briefly in 1x phosphate-buffered saline, destained twice for 30 min with phenol-red free DMEM, and transferred to fresh phenol-red free DMEM (Gibco #31053036).

For OLS imaging, an automated script was used to acquire data across multiple fields of view per condition. Receiver fluorophores (JFX650) were shelved by illuminating the sample with 642 nm light at full power (2.2 W at the laser head). Movies were recorded under continued 642 nm illumination at a frame rate of 10 ms per frame. After acquiring 50 baseline frames, the sample was illuminated with either 405 nm light at full power (100 mW at the laser head) or 561 nm light at 50% AOTF (1.1 W at the laser head) for 50 frames, and 50 more frames were imaged to detect reactivated fluorophores. Receiver dyes were then re-shelved using a second pulse of 642 nm light for 50 frames, and this sequence was repeated for three cycles, alternating between 405 nm (direct reactivation) and 561 nm (PAPA).

Laser output through the objective was measured prior to the experiment using a slide-mounted power sensor (Thorlabs S170C) at levels compatible with the sensor’s detection range. Under these conditions, measured powers were 0.65 mW for 405 nm (full power), 22 mW for 561 nm (5% AOTF), and 30 mW for 640 nm (200 mW laser head power). These measurements served to validate the consistency of laser output across sessions. To verify measurement reproducibility, a positive control (Halo-Sp1 + SNAP-Sp1) and negative control (Halo-Sp1 + SNAP-NLS) were imaged at the beginning and end of each imaging session.

Klf1-expressing cells were imaged using highly inclined and laminated optical sheet (HILO) illumination on a custom microscope previously described by Hansen et al. (2018). Approximate laser power densities at the sample were 52 W/cm^2^ for 405 nm (violet), 100 W/cm^2^ for 561 nm (green), and 2.3 kW/cm^2^ for 639 nm (red). Emission was collected through a Semrock 676/37 bandpass filter.

### PAPA-SMT data analysis

Molecules were tracked using quot (https://github.com/alecheckert/quot), a Python toolkit for high-accuracy single-molecule localization and trajectory analysis. Nuclei were segmented using the Cellpose algorithm (Stringer et al., 2021), applied to 560 nm fluorescence images after Gaussian smoothing (σ = 10), with an estimated nuclear diameter of 80 pixels. Segmented regions smaller than 5,000 pixels in area were excluded, and adjacent or touching nuclei were separated using a custom boundary-drawing routine. Nuclear masks were overlaid on 642 nm intensity images to compute a per-nucleus leakage score, defined as the mean 642 nm signal in a 4-pixel ring surrounding the nuclear mask divided by the nuclear mean intensity. Cells with high leakage scores (>0.7) were excluded to eliminate those exhibiting cytoplasmic 642 nm signal. Additional filters excluded nuclei with atypically low or high detection counts and those contacting the top or bottom edge of the image field. Trajectory data were intersected with nuclear masks to assign detections to individual cells, and only cells passing all criteria were retained for analysis. Diffusion coefficient distributions were inferred using SASPT (https://github.com/alecheckert/saspt), a Bayesian state array framework for modeling heterogeneous mobility states from single-molecule trajectories.

To calculate proximity indices, detections were counted in each frame within segmented nuclear masks and grouped into four time windows per cycle: frames 0–49 (baseline preceding violet pulse), 50–99 (violet-induced direct reactivation, DR), 100–149 (baseline preceding green pulse), and 150–199 (green-induced proximity-assisted photoactivation, PAPA). To correct for spontaneous reactivation of dark-state Janelia Fluor dyes—which occurs independently of laser input and follows an exponential decay—baseline counts were fit to a single-exponential model and extrapolated into the corresponding post-pulse window. This projected background was subtracted from the observed detections, and negative values were truncated to zero. The total number of background-corrected green detections (PAPA) was divided by the number of background-corrected violet detections (DR) to yield a green-to-violet (GV) ratio. Each experimental GV ratio was normalized to the GV ratio of a non-interacting SNAPf-3xNLS negative control to compute the final proximity index. A proximity index of 1 indicates no increase in proximity-induced signal above background. To estimate statistical uncertainty, GV ratios were recalculated using bootstrap resampling of individual cells (100 replicates per condition). Error bars represent the 2.5th and 97.5th percentiles of the resulting bootstrap distribution. Uncertainty in the control condition is shown as a fixed error bar at a proximity index of 1.

To visualize the relationship between DR and PAPA-reactivated molecules at the single-cell level, we plotted background-corrected PAPA detections (green channel) against DR detections (violet channel) for each cell across three imaging cycles. For each pulse type, background was estimated as the number of detections in the 50 frames preceding the pulse, and subtracted from the number observed in the 50 frames following the pulse. Cells with low detection counts (<100) or negative corrected values were excluded. A linear regression was performed without intercept to quantify the proportionality between green and violet signals, and 95% confidence intervals for the fitted line were estimated by bootstrapping cell samples (500 replicates).

### Fluorescence recovery after photobleaching (FRAP)

FRAP experiments were performed using a Zeiss LSM 900 confocal microscope. Cells labeled with fluorescent dye were imaged using a 561 nm laser and a 150 µm pinhole. Imaging was conducted at the maximum scan speed permitted by the frame size, with bidirectional scanning and 8-bit depth. Image sequences were acquired with a 150 ms interval between frames for a total of 600 frames (90 seconds). Photobleaching was initiated at frame 16, following a 15-frame pre-bleach baseline period used for normalization. 20 movies were acquired per condition, and each condition was imaged in two independent experiments conducted on separate days.

### FRAP data analysis

FRAP movies were analyzed using a custom Python pipeline (https://github.com/vinsfan368/FRAPanalysis). A still from each movie was used to generate a nuclear mask, which was smoothed and thresholded to segment the nucleus. The position of the bleach spot was extracted from CZI file metadata and corrected for lateral drift using 2D image cross-correlation.

For each frame, the mean fluorescence intensity within the bleach spot was measured. Background signal was estimated as the median pixel intensity outside the nuclear mask. Recovery was calculated as the background-subtracted intensity ratio of the bleach spot to the full nucleus. This ratio was normalized to the average ratio over the pre-bleach frames (frames 1–15), such that full recovery corresponds to a normalized value of 1.

### Western blotting

Cells were lysed directly in SDS sample buffer by pelleting approximately 1 × 10^6^ cells from a six-well plate, resuspending in 100 µL of SDS buffer, and heating at 95 °C for 10 minutes.

Lysates were resolved by SDS-PAGE and transferred to nitrocellulose membranes using wet transfer at a constant current of 90 mA overnight in transfer buffer (15 mM Tris-HCl, 20 mM glycine, 20% methanol, 0.0375% SDS). Membranes were briefly stained with Ponceau S to verify transfer, then blocked for 1 hour at room temperature in TBS containing 5% bovine serum albumin (BSA) with gentle agitation. Primary antibody incubation was carried out overnight at 4 °C in TBST containing 5% BSA. To detect Sp1, membranes were incubated with anti-Sp1 mouse monoclonal antibody (Santa Cruz Biotechnology, sc-17824). After three washes in TBST, membranes were incubated for 1 hour at room temperature with an HRP-conjugated anti-mouse secondary antibody (1:5000 dilution in 5% BSA in TBST). Blots were then washed three times in TBST with brief (∼2-minute) incubations under agitation prior to chemiluminescent detection.

## Acknowledgments

Thanks to Eric Betzig for sharing a home-built OLS microscope design, Amir Hay, Ziyuan Chen, and Gokul Upadhyayula for help with microscope construction and troubleshooting, and Dan Milkie for assistance with software. This work was supported by early-stage investigator developmental funds from NIH grant RM1GM139738 (to T.G.). R.T. is an Investigator of the Howard Hughes Medical Institute. We would like to thank the Tjian-Darzacq group, the members of our RM1 collaboration team, as well as Aditya Udupa, Connor Horton, Max Staller, Joel Auerbach, Ben Knepper, Izaak Meckler, and Adam Hochschild for helpful discussions.

## Notes

### Competing Interest Statement

The authors have declared no competing interest.

### Summary of Updates

We corrected typographical errors, improved clarity and organization of the text, integrated high-resolution figures into the main manuscript, and expanded the reference list. No substantive changes were made to the results or conclusions.

